# Computer simulations reveal mechanisms of spatio-temporal regulation of DNA replication

**DOI:** 10.1101/2024.05.24.595841

**Authors:** Razie Yousefi, Maga Rowicka

## Abstract

The dynamics of DNA replication, activation of replication origins and elongation rate, depend on dNTP in ways that remain poorly understood. Here, we present RepliSim, a probabilistic computer model of DNA replication simulation, which is capable of simulating DNA replication not only in normal conditions, but also during perturbations (e.g. HU treatment) and in mutant cells (here checkpoint deficient yeast cells). We show that different mutations in checkpoint genes, affect the origin activation differently by modulating the stochasticity of firing time and efficiency, regardless of the dNTP levels. The origin activation is less affected by increase in dNTP levels, however it declined drastically during the S phase when the dNTP levels decreased compared to G1 phase. Additionally, we show that the distribution of distances covered by replication forks is influenced not only by average fork progression speed, but also by firing time and fork progression speed stochasticity, while the latter has been neglected in most of the proposed models for DNA replication. These together, opens up new insight to the studies about the dynamics of replication and how it is coordinated during the S-phase.

## Introduction

DNA replication in Eukaryotic cells is highly regulated to ensure that the whole genome is duplicated correct, complete, and only once in a timely manner [1]. DNA replication initiates at specific cites, termed origins of replication, which are determined and prepared to fire with the assembly of pre-replication complex (pre-RC) during late M-phase and early G1-phase [2]. Replication origins are licensed in exes and during the subsequent S-phase a subset of origins initiate replication from which two forks are emanated and elongate bi-directionally [3]. In the budding yeast *Saccharomyces cerevisiae*, DNA replication initiates from hundred of well defined, site specific origins and their activation is partly in a chronological programmed order and partly stochastically [4]. As a result, even in the same cell population, the choice of origins to be activated varies from cell to cell. This flexibility in the active origin population is essential in the response to DNA damage and adaption of replication to gene expression and cell fidelity, however it has not been yet determined how origins are selected to be activated [5].

To execute replication, deoxyribonucleotids (deoxynucleoside triphosphates [dNTPs]) are required by cells during the S-phase. As dNTP pools present in G1 are not sufficient to replicate the whole genome, it is important for ribonucleotide reductase (RNR) activity (the only mechanism for synthesis of deoxyribonucleotids [5]) to be maintained and highly regulated to coordinate with the DNA synthesis through out the S-phase [6].

It has been shown that alteration of dNTP levels can affect the replication fork speed, for example, DNA replication is slowed down in the presence of hydroxyurea (HU), which partially depletes nucleotide precursor pools by inhibiting RNR activity and as a result preventing the reduction of ribonucleotides to dNTPs [7,8]. Further studies of how dNTP levels impact the replication progression and fork speed, will help us better understand the dynamics of DNA replication.

To ensure the accurate replication, yeast cells have developed complex surveillance mechanisms called checkpoints that monitor the DNA replication, detect and and stabilize the stalled forks, and regulate the origin activation and fork elongation by alteration of the cellular metabolism to avoid genetic instability following replication stress [9].

Apart from expansion and consumption of dNTP pools during replication, DNA lesions also impact the dNTP levels. It has been suggested that in the presence of DNA lesions, expansion of dNTP pools, by activation of checkpoints, helps cells to promote S-phase progression [10,11]. Checkpoint pathways ensure the proper order and timing of cell cycle activities. To study the checkpoint response to HU, it is important to understand how checkpoint dependent mechanisms regulate dNTP pools in order to promote integrity. It has been shown that checkpoint response to HU treated cells is dependent on the protein kinases mec1/rad53 pathways [12,13,14]. mec1 is required to activate checkpoints including rad53, which regulates the cellular responses in the presence of genotoxic agents [15,16]. It is also necessary for phosphorylation and diminution of sml1, a gene whose deletion is essential for the viability of mec1Δ and rad53Δ mutants [17,18].

In different checkpoint deficient cells, different levels of dNTPs are observed, in which replication rate is proportional to the dNTP level [6]. For instance, during late G1 and early S-phase, dNTP level in HU treated mec1-1sml1-1 (mec1) mutant cells is multiple times larger than that in wild type (wt) cells. From the other side, fork progression studies in HU treated wt and mec1 cells show that BrdU tracks are significantly longer in mec1 compared to the wt cells. This suggests a larger fork progression rate in HU treated mec1 cells, which is due to a higher dNTP level.

It has been claimed that alteration of dNTP pools by dysregulation of RNR activity affects the origin activation [5,6]. During HU-induced stress, late origins are specifically inactivated in wt cells so that the observed ssDNA correspond to early firing and/or high efficient origins [19]. In addition, fork elongation is slowed down significantly in the presence of HU compared to unchallenged forks. However, in budding yeast cells exposed to HU are able to complete the S-phase, though it is slowed down considerably compared to the untreated cell cycle (90-120 min) [5,9].

DNA replication in HU treated cells has been studied widely in wt as well as checkpoint deficient cells. In HU treated wt cells, replication corresponds to activation of a subset (early/more efficient) of origins, while in mec1 mutant cells for instance, most of the potential origins are active as expected since rad53, which modulates the temporal program of firing of origins by repressing late origins, is inactivated as a result of mec1 mutation [20].

However, it is still unclear how alteration of the dNTP level in response to HU impacts the mechanism of origin activation in different conditions and experimental data is not able to unmask all the details regarding to the dynamics of replication corresponding to each individual condition.

In this work, we present a porobabilistic numerical model for DNA replication simulation (RepliSim), which examines replication in the HU induced wt as well as checkpoint deficient (mec1, mrc1Δ, mrc1^_AQ_^, and rad53-11(rad53), rtt101Δ, ctf18Δ, rad52Δ, sgs1Δ, asf1Δ, elg1Δ, rrm3Δ, and ctf4Δ) cells. Our model includes defined origin position, probabilistic initiation time and fork elongation rates assigned to origins and forks using a MonteCarlo method, and a transition time during the S-phase at which origins transit to a silent/non-active mode from being active. The model and parameter selection is done by fitting it to the microarray and fiber experimental data available in [6].

We show that expansion of dNTP pools does not impact the origin activation, however at certain point during the S-phase origins of replication enter a silent/non-active mode, when the dNTP pools is at a certain relative level compared to that at G1. This transition could be a result of the limited number of origin initiation activation factors as well. Additionally, simulations reveal that different checkpoint deficiencies affect the firing time of origins by influencing the stochasticity of firing time. As stochasticity of firing time increases, origins including late ones, get a higher chance to fire simultaneously. This causes the observation of many active origins, including late origins, in some of the conditions.

Moreover, simulations reveal that variation in the fork elongation speed affects the distribution of the distances covered by replication forks, while fork progression speed has been assumed to be constant in most of the proposed models and simulations investigating the dynamics of DNA replication.

In our simulation experimental data is utilized for modeling and validation of our results.

## Results

In this study, we are interested in understanding the mechanism of DNA replication in HU treated wt and checkpoint deficient cells and investigating how the kinetic of replication is associated with the dNTP levels. To address this question, we developed a probabilistic numerical method for DNA replication. The experimental data used for the simulation includes DNA fiber data, microarray-based data, and dNTP pools levels available in [6], which indicates how dNTP levels alter the execution of DNA replication.

### Mutants impact origin activation

Replication profile experimental data conveys a high-density of active origins in mec1, mec1-100, rad53, mrc1Δ, and ctf18Δ mutant compared to the wt, mrc1^_AQ_^, rtt101Δ, rad52Δ, sgs1Δ, elg1Δ, rrm3Δ, and ctf4Δ cells while, the amount of dNTP levels is very different in these mutations as shown in Fig 1(a). This indicates that there is not an obvious correlation between dNTP levels and density of active origins. For instance, in wt cells about 300 origins are active and in rad53 the number of active origins are more than 400, while in both cells the amount of the dNTP pools is relatively similar both in G1 and 60 minutes from S-phase.

**Fig 1.**
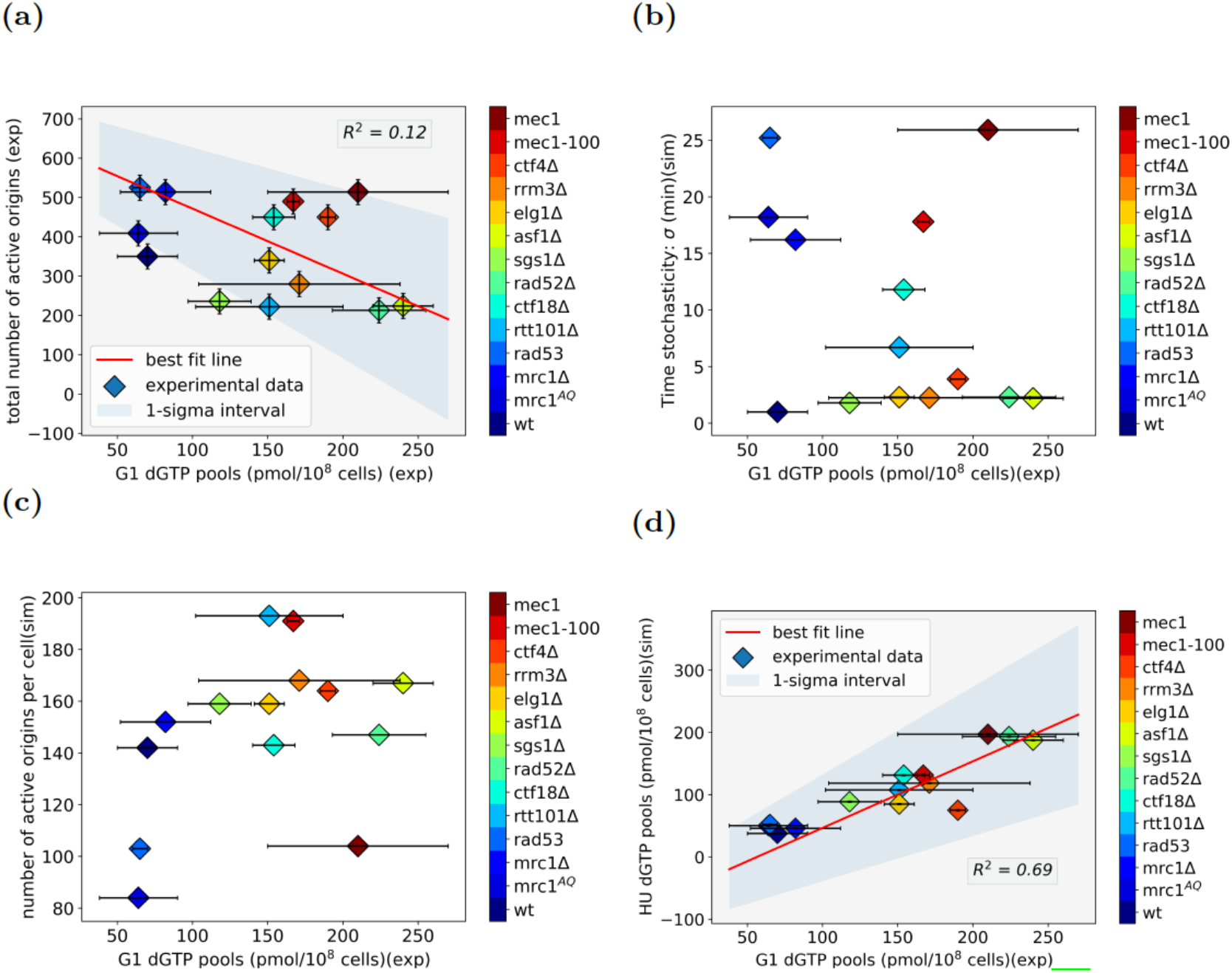
A representation of observations based on experimental data in [12] and simulation results in the indicated cells exposed to HU. (a) Scatter plot of experimental results for both total number of active origins observed from BrdU-IP chip analysis versus the amount of dNTP level. As shown in the figure, there is not an obvious correlation between the origin activity and the amount of dNTP pools in the cell environment (R_2_ = 0:12). (b) A representation of the stochasticity of firing time (*σ*_*t*_) derived from simulations and the experimental G1 dNTP pools level, which indicates that the firing time randomness is independent of the amount of dNTP pools. (c) The total number of active origins per cell predicted from simulation versus the dNTP pools at G1 from experiment represents no obvious correlation between origin activity and the amount of building blocks for DNA synthesis. (d) Schematic representation of dNTP levels at t___ (time from G1 when origins of replication enter a silent/non-active mode) derived from simulations and its experimental value at G1, which shows that origin activity can be affected by the relative amount of dNTP pools.

In addition, most of the potential origins, including late origins, get the chance to fire almost simultaneously in high-density active origin cells, while in the rest of the conditions, the number of active origins increases in a gradual fashion. Simulations reveal that in HU treated high-density active origin cells, the firing time of origins are much more stochastic compared to the normal replication (Fig 1(b)). For instance, in HU induced mec1 cells, the stochasticity of firing time of origins is about four times higher than wt cells. As randomness of the firing time increases, late origins get a higher chance to fire during the early of S-phase, which causes the observation of many active origins including late origins. This means that mec1, mec1-100, rad53, mrc1Δ, and ctf18Δ mutations, directly or indirectly, disturb the temporal program of origin firing by making them more random while, the mean firing time is relatively in the same order as in wt cells. This finding is consistent with previous observations where, it has been shown that mec1 and rad53 modulate the firing time of origins, but unclear how exactly controls the temporal program of replication [20].

It has been shown that certain mutations cause chromosome condensation and compaction [21,22]. Considering this finding, one plausible explanation is that specific mutations make the DNA molecule less compact and as a result, facilitate the access of the replication initiation factors to the origins of replication including late origins. Another evidence which makes this explanation more feasible, is DNA replication of HU treated wt cells at two different temperatures, 25 and 30°c, in which at higher temperature less number of active origins has been observed [6]. From the other side, at higher temperature DNA molecule shows a more compact folding structure [23]. This is in agreement with our postulation. Furthermore, it has been shown that placing early active origins near a telomere, forces it to fire later in the S-phase [24], which indicates that the firing time of origins is not an intrinsic property of specific origin sequences and could be related to the topology of DNA molecule.

Still, there is a difference between the density of active origins in cell population and the number of active origins per cell, where the former can be observed from experimental data, while the latter can be predicted by simulations. As shown in Fig 1(c), from simulations the number of active origins per cell is also independent of the amount of dNTP pools and is modulated by mutations.

Moreover, simulations reveal that during the S-phase, regardless of being early or late, origins of replication make a transition from being active to a silent/non-active mode where probability of origins to fire decreases drastically to less than 0.0001 of the competence of each origin, while replication continues with an average steady rate consistent with [5]. This transition occurs in most of the cases when dNTP pools drop down to a certain level relative to the dNTP pools measured at G1 as shown in Fig 1(d). Though, this transition could be a result of the limited number of replication initiation factors available in the cell environment or some factors limiting the number of replication forks.

### Extended dNTP pools promotes fork elongation

Analysis of DNA content confirms a slower fork progression in HU treated cells compared to the untreated cells yet, in the presence of HU, different mutations have different fork progression rates since they modulate the amount of the dNTP polls differently. Experimental data reveals an overall longer distances covered by forks for larger dNTP pools since expansion of dNTP pools delays the activation of checkpoints that inhibit fork elongation. Analogously, as shown in Fig 2(a), from simulations a higher fork progression speed is detected for conditions with larger dNTP pools, which causes a longer distance covered by individual forks, however dNTP level is not the only factor playing a role in the replication rate, since as shown in Fig 2(b), some of the mutations, including asf1Δ, rad52, and mec1, are among the cells containing the largest amount of the dNTP pools at G1 while their fork speed is not the highest. This shows that other than dNTP level, other mechanisms also affect the replication rate in different mutations. In Fig 2(c), all the cell conditions are categorized in three different groups considering their relative replication rate, which shows that more biological investigations are needed to examine characteristics of different mutations and their affect on the replication rate.

**Fig 2.**
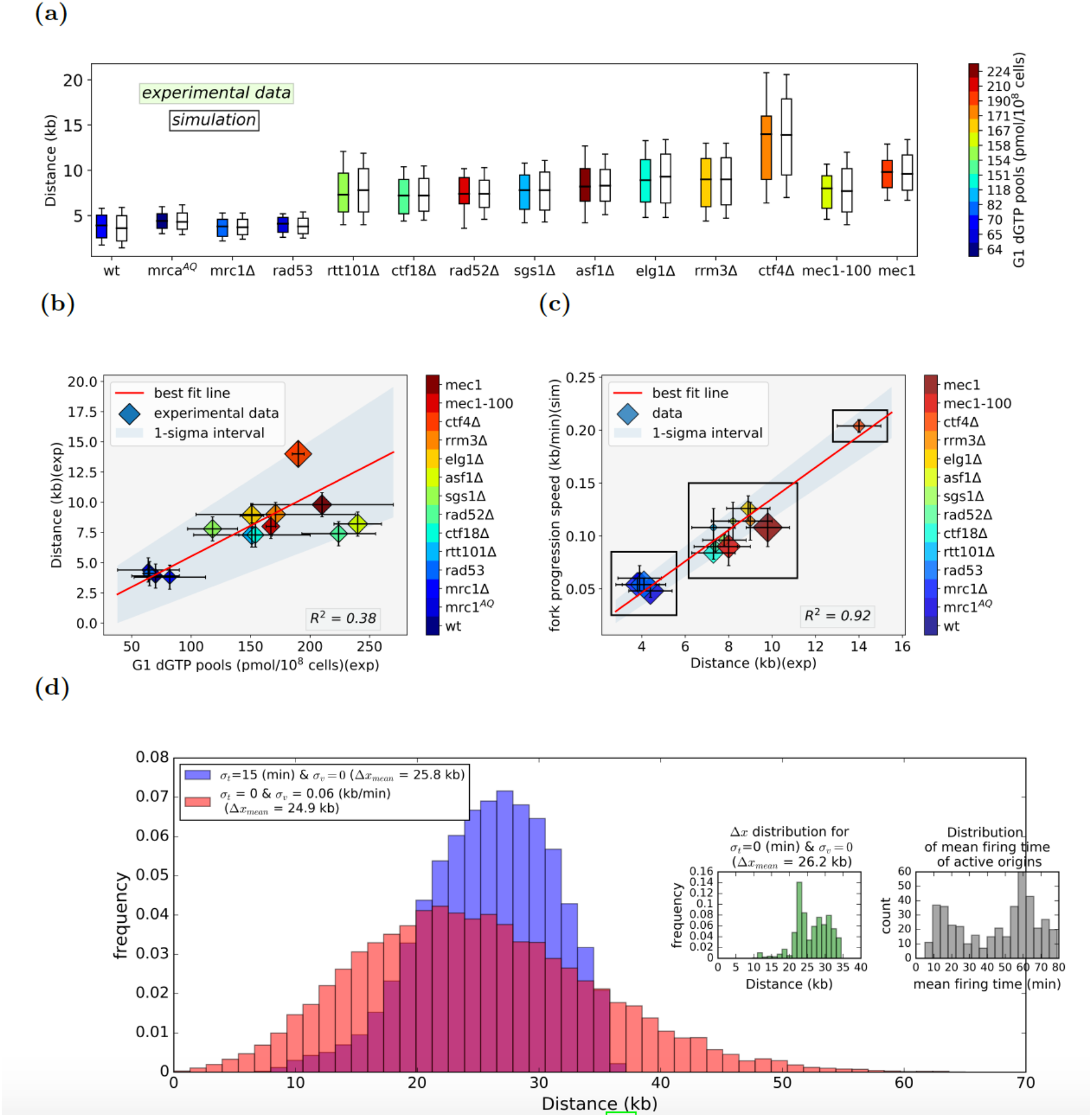
A representation of experimental data in [12] and simulation results in different HU treated cell conditions. (a) A comparison between experimental data and simulation results for the length of the replicated DNA. Box and whiskers indicate 25-50-75 and 10-90 percentiles, respectively. As shown, there is a good agreement between experimental data and simulation results. (b) An illustration of the experimental results for G1 dNTP pools and the mean distance covered by individual forks which shows that a larger amount of dNTP pools leads to a higher fork progression speed, however the correlation is not very strong (R_2_ = 0:39). Stochasticity of firing time is shown as the area of the markers in the scatter plot. (c) Scatter plot of fork progression speed versus mean distance covered by replication forks, obviously classifies different cell conditions to three groups. Error bars for speed represent standard deviation for each individual condition.(d) The distribution of _x for highly stochastic firing time, highly variable fork speed progression, and constant speed-non-variable firing time from simulations for a random set of parameters. Noticeably, the distributions are different in all three cases. For a better illustration, constant speed and non-variable firing time distributions are presented on the same axis, which demonstrates the difference between them very well. Additionally, the distribution of constant speed&non-variable firing time and mean firing time of active origins are presented side by side and as expected, they are in the same fashion. The average distance traveled by replication forks (Δ*x*_mean_) is comparable in all three cases.

A longer replicated distance could be a consequence of a longer time for traveling forks (which is equivalent to an earlier firing time of origins during S-phase), a larger fork speed, or a combination of both. This is one of the questions we address here. From both simulations and analysis (Equ. (6)), the average distance traveled by replication forks (Δx_mean_) depends on the mean firing time of the origins and mean fork speed, however the variance (stochasticity) of the distribution of Δx is affected by the variance of both firing time of origins and fork progression speed as well.

Fig 2(d) represents a comparison of three different cases: a highly stochastic firing time/ constant speed (*σ*_v_ = 0), a highly stochastic fork speed/firing time specific (*σ*_t_ = 0), and constant speed & firing time specific (*σ*_v_ = *σ*_t_ = 0 = 0). As shown in the figure, Δx_mean_ in all three cases are comparable confirming the assumptions we have made to build up our model. While, there is a significant difference between Δx distributions for a variable fork speed compared to a constant speed. This is in contrast with the assumptions which have been made in previous proposed computational analysis and models, which investigate the dynamics of DNA replication [25,26,27,28,29], where in most of the cases the comparison between proposed models and experimental data, has been made between individual origins and not genome wide. In addition, in most of the proposed models, the average of different quantities played a role in model selection, which as shown in here, could be miss-leading. This result shows that considering a constant fork progression speed is not an accurate assumption.

### Validation of results from simulations

In order to validate our model and parameter selection, we compared simulation results for the amount of the replicated genome after 60 min from the S-phase with that of the experimental data. As shown in Fig 3(a), for most of the conditions, the results are quite compatible. In rad52Δ, mec1-100, and mec1 mutants, there is a difference between our predictions from simulation and experimental data, where simulations reveal that the difference in the dNTP pools is less than what we observed from experiment. This could be an error in the experimental data collection or some mechanism inside the cell, which consumes the dNTP pools such as transcription and gene expression, which is left for further investigations by biologists. It is noticeable that all of these mutant cells are among the cells with higher replication rate.

**Fig 3.**
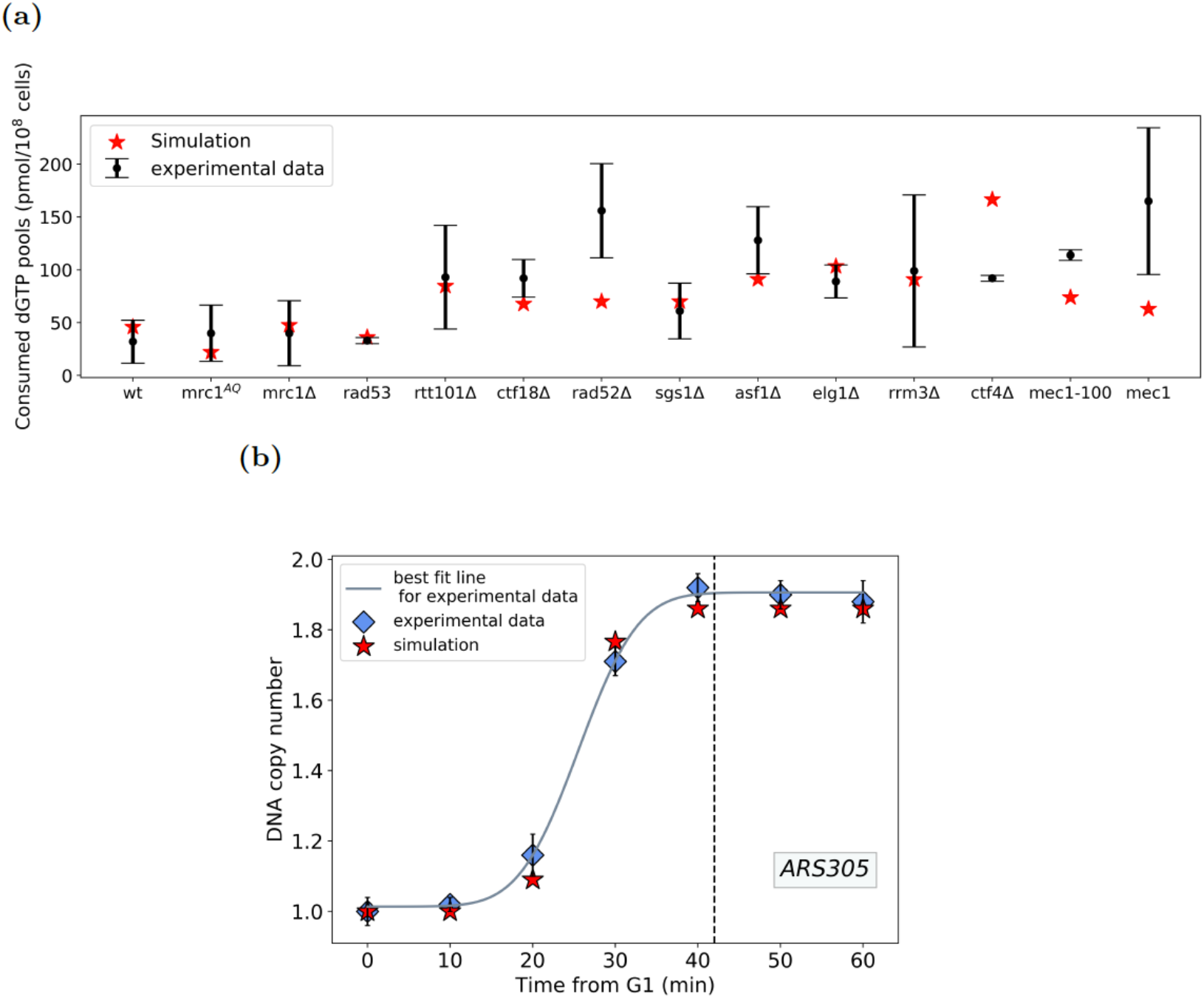
An illustration of experimental data in [12] and simulation results. (a) A comparison of the amount of consumed dNTP pools from simulation and experimental data. The results are in agreement for most of the cases, however in some of the mutations (rad52_, mec1-100, and mec1) a smaller amount of consumed dNTP pools is predicted, which could be an experimental error or some unknown mechanism which consumed the dNTP pools more that other conditions. (b) A schematic of the kinetics of ARS305 loci from both experimental data and simulations, which illustrates their compatibility. As shown, in both cases the DNA copy number remains almost constant after _40 min from G1, that confirms our model based on which during S-phase origins of replication transit from an active mode to a silent/non-active mode.

Another value from simulation for HU treated wt cells is the transition time, when during S-phase origins of replication transit from an active mode to a silent/non-active mode, is derived to be equal to 42.6 minutes. Interestingly this time is in complete agreement with the BrdU micro-array DNA copy number of ARS305, an efKicient early origin, where after 40 minutes from S-phase, the DNA copy number remains the same meaning a negligible origin activation at this loci. Fig 3(b) represents the kinetics of ARS305 loci from both experimental data and simulations.

## Materials and methods

### Simulations

We developed a probabilistic numerical model which takes in to account two groups of parameters; local and global. Local parameters are individual to each specific origin, while global parameters are those, which are assumed to be approximately similar all over the genome. To build up the model, we made the following assumptions:

1. Cells are independent and individual chromosomes of each cell are also independent.
2. Origins of replication are site determined and their coordinates along the chromosomes are verified and available experimentally.
3. A probability of licensing *p*_*i*_ (competence) is assigned to each individual origin based on which an origin is biochemically ready to fire. *p*_*i*_ is related to the (experimentally measured) efficiency of the origins (*i*) as *e*_*i*_ ≤ *p*_*i*_ ≤ 1, where *e*_*i*_ = *n*_*i*_ /*N*, with *n*_*i*_ as the total number of cells in which th origin is replicated actively and *N* as the total number of cells.
4. Firing time of replication for each individual origin is *t*_*i*_, which is generated using a Monte-Carlo method from a Gaussian probability distribution with an experimentally estimated mean firing time 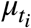 specific to that origin, and a global standard deviation α*t*;

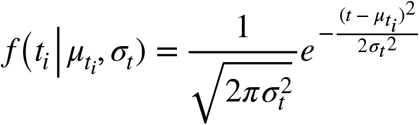
5. Individual forks progress with different speeds, generated using a Monte-Carlo method from a Gaussian probability distribution with a global average speed *μ*_*v*_ and standard deviation *σ*_*v*_, and is maintained uniform during the progression;

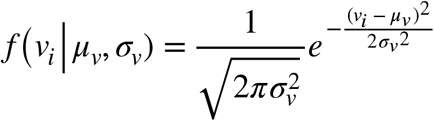
6. During the S-phase, the probability of origin firing can change with a factor after a duration t_α,_ when (some) mechanisms in the cell inhibit more origin activation.

We used a list of 529 origins detected in [6] with the experimentally derived efficiencies assigned to each origin. These origins are compatible with the OriDB database [30]. We used a Monte-Carlo method to assign a firing time and two speeds (for the two forks emanating from that origin) to each individual origin with a probability *p*^*i*^ (*e*_*i*_ ≤ *p*_*i*_ ≤ 1). Using these initial parameters, our model is able to take into account the passive replication by selecting only active firing origins. Then the replicated distances from each individual actively fired origin are derived and used to compare with that of the experimental data for model and parameter selections .The whole genome experimental data for distribution of distances covered by forks emanating from the same origin are utilized by minimizing the sum of the square differences between experimental data and simulation. For minimization, five percentiles, 10, 25, 50, 75, and 90^th^, of both experimental and simulation data are used to calculate the sum of square differences.

A genetic algorithm is used for minimization of the sum of the square of differences between experimental data and simulation. We used a population of 10000 sets of parameters, run in parallel using the open source implementation OpenMP over 32 threads. For each condition, a number of best sets of parameters is selected among which the one is chosen with more similar total number of active origins compared to the experimental data, as shown in Fig 4.

**Fig 4.**
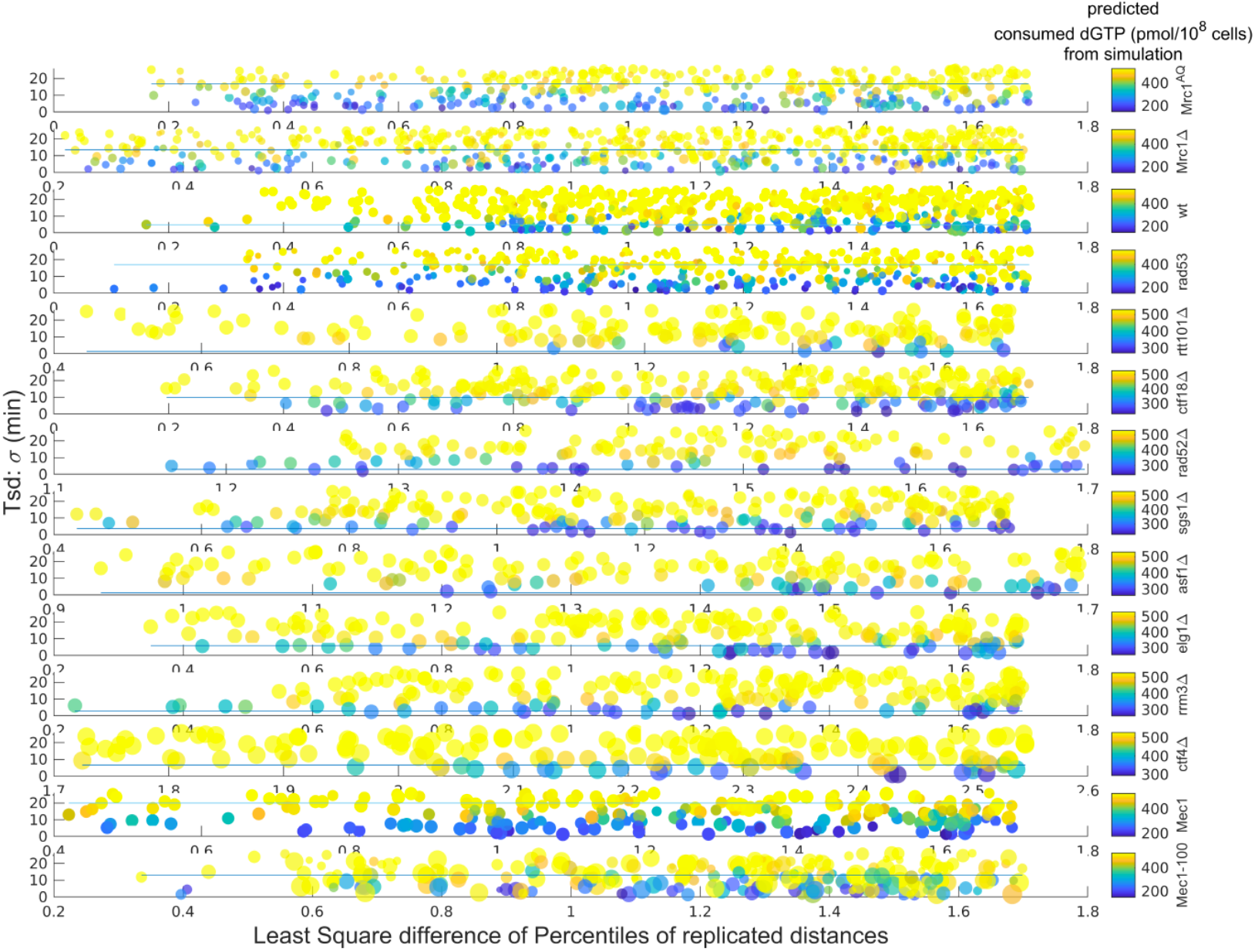
Simulation results for different models with the set of parameters best in agreement with the experimental data. The color-bars represents the total number of active origins which are larger in higher stochastic firing time (y axis). Among these, a model is chosen with more similar number of total active origins compared to that from experimental data, shown with a horizontal line in each condition.

### Mean firing time from experimental data

Mean firing time of individual origins is determined from BrdU-labeled microarray experimental data available in [c] where at each individual origin, the distribution of the tracks (Δ*x*) of the replicated DNA, originated from the origins of replication, is calculated and used to derive the mean firing time considering the following assumptions;

1. To normalize the distribution of DNA tracks measured from BrdU experimental data, we used the BrdU micro-array DNA copy number of *ARS305* and normalized it to give the same efficiency as derived from its DNA copy number from quantitative PCR experiment.
2. In the presence of HU, mean firing time of origins in mutant cells remains in the same relative order as in wt cells, based on the observations in 12].
3. Mean firing time of each origin is individual to that origin, however variation of firing time from the mean (*σ*_*t*_) is the same for all the origins and assumed to be a global parameter. Then, the firing time of each origin, which is also assumed as the initial time of firing (*t*_0*i*_), is derived from the following normal distribution:

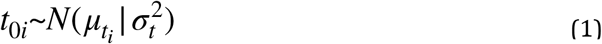
4. Individual forks have different speeds, however the speed of each fork is derived from the same probability distribution with a mean speed equivalent to that observed from experiment:

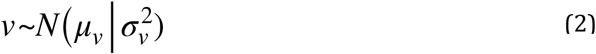

Considering the relation Δ*x* = *v*. Δ*t*, knowing the distribution function of Δ*x*, we are able to derive the distribution function of the firing time for each origin from which the mean firing time can be predicted as follows:

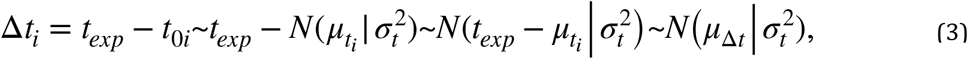

Where 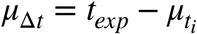.

From the other side, assuming *σ*_*t*_*σ*_*v*_ ≪ *μ*_Δ*t*_ *μ*_*v*_ [31], we have

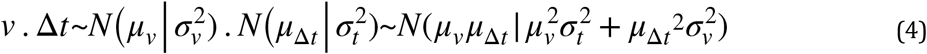

Considering Equ. (4) and taking into account the assumption 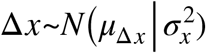 together, we have:

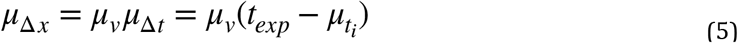

from which we derive the mean firing time of each individual origin as

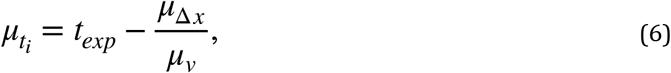

which is used in our simulations to infer the mean firing time by implementing *t*_*exp*_ and *μ*_Δ*x*_ while *μ*_*v*_ is the parameter, which is adjusted in the simulation through parameter selection in the genetic algorithm.

## Conclusion

Here, we present a probabilistic numerical model for DNA replication simulation (RepliSim), which examines the dynamics of replication in HU treated wt as well as checkpoint deficient (mec1, mrc1Δ, mrc1^AQ^, and rad53, rtt101Δ, ctf18Δ, rad52Δ, sgs1Δ, asf1Δ, elg1Δ, rrm3Δ, and ctf4Δ) yeast cells. We utilize experimental data for model and parameters selection and validation of our results.

Experimental data reveals that expansion of dNTP pools does not impact the origin activation. Concurrently from simulations the firing time of origins’ stochasticity is different in various mutant cells. A higher stochasticity in firing time gives origins including late ones, a higher chance to fire almost simultaneously. Consequently, in some of the checkpoint deficient cells many active origins are observed. As DNA molecule compaction is affected in the presence of different mutations [32,33], origin replication initiation factors get a higher chance to access origins and give them a higher chance to fire earlier in time. Another evidence which makes this assumption more feasible is DNA replication of wt yeast cells in the presence of HU at 25 and 30°c, where a more compact DNA molecule is observed at higher temperatures and proportionally, at 30°c less number of active origins is observed.

Additionally, from simulations, origins of replication make a transition from being active to a silent/non-active mode, when probability of origins to fire declines drastically. This transition occurs in most of the cases when dNTP pools drop down to a certain level relative to the dNTP pools measured at G1, however it could be a result of the limited number of replication initiation factors available in the cell environment.

Both experimental data and simulations propose that expansion of dNTP pools promote fork progression and as a result observation of longer replicated distances. Although, in the presence of various mutations the fork progression rate is modulated differently. Simulations and analysis reveal that the variance of distribution of the distances covered by replication forks is not only dependent to the fork progression speed but also randomness of fork speed and firing time as well as initial firing time of each specific origin. This finding is in contrast with the proposed replication models in which fork progression rate is assumed to be constant.

DNA replication and replication origin activation are intricately linked to chromatin structure, which modulates the accessibility and efficiency of replication origins. Chromatin remodeling and histone modifications play pivotal roles in ensuring precise initiation and progression of DNA replication. Future research should focus on elucidating the mechanisms by which chromatin dynamics influence origin activation and the mechanism of DNA replication. Additionally, improving RepliSim for application to the human genome, which remains un-modeled due to its complexity, will enhance our understanding of DNA replication mechanisms. These areas will be the focus of our future research.

## Authors Contributions

Razie Yousefi, Formal analysis, Investigation, Methodology, Software, Validation, Writing – original draft, Writing – review & editing and Maga Rowicka, Conceptualization, Funding acquisition, Investigation, Methodology, Supervision, Writing – original draft, Writing – review & editing.

